# Host tracheal and intestinal microbiomes inhibit *Coccidioides* growth *in vitro*

**DOI:** 10.1101/2023.10.23.563655

**Authors:** Susana Tejeda-Garibay, Lihong Zhao, Nicholas R. Hum, Maria Pimentel, Anh L. Diep, Beheshta Amiri, Suzanne S. Sindi, Dina R. Weilhammer, Gabriela G. Loots, Katrina K. Hoyer

## Abstract

Coccidioidomycosis, also known as Valley fever, is a disease caused by the fungal pathogen *Coccidioides*. Unfortunately, patients are often misdiagnosed with bacterial pneumonia leading to inappropriate antibiotic treatment. Soil bacteria *B. subtilis*-like species exhibits antagonistic properties against *Coccidioides in vitro*; however, the antagonistic capabilities of host microbiota against *Coccidioides* are unexplored. We sought to examine the potential of the tracheal and intestinal microbiomes to inhibit the growth of *Coccidioides in vitro*. We hypothesized that an uninterrupted lawn of microbiota obtained from antibiotic-free mice would inhibit the growth of *Coccidioides* while partial *in vitro* depletion through antibiotic disk diffusion assays would allow a niche for fungal growth. We observed that the microbiota grown on 2xGYE (GYE) and CNA w/ 5% sheep’s blood agar (5%SB-CNA) inhibited the growth of *Coccidioides*, but that grown on chocolate agar does not. Partial depletion of the microbiota through antibiotic disk diffusion revealed that microbiota depletion leads to diminished inhibition and comparable growth of *Coccidioides* growth to controls. To characterize the bacteria grown and narrow down potential candidates contributing to the inhibition of *Coccidioides*, 16s rRNA sequencing of tracheal and intestinal agar cultures and murine lung extracts was performed. The identity of host bacteria that may be responsible for this inhibition was revealed. The results of this study demonstrate the potential of the host microbiota to inhibit the growth of *Coccidioides in vitro* and suggest that an altered microbiome through antibiotic treatment could negatively impact effective fungal clearance and allow a niche for fungal growth *in vivo*.

**Importance:** Coccidioidomycosis is caused by a fungal pathogen that invades host lungs, causing respiratory distress. In 2019, 20,003 cases of Valley fever were reported to the CDC. However, this number likely vastly underrepresents the true number of Valley fever cases as many go undetected due to poor testing strategies and lack of diagnostic models. Valley fever is also often misdiagnosed as bacterial pneumonia, resulting in 60-80% of patients being treated with antibiotics prior to accurate diagnosis. Misdiagnosis contributes to a growing problem of antibiotic resistance and antibiotic induced microbiome dysbiosis, and the implications on disease outcome are currently unknown. 5%-10% of symptomatic Valley fever patients develop disseminated and/or chronic disease. Valley fever causes a significant financial burden and reduced quality of life. Little is known regarding what factors contribute to the development of chronic infection and treatments for disease are limited.

## Introduction

*Coccidioides immitis* and *Coccidioides posadasii* are soil fungi responsible for the disease coccidioidomycosis, also known as Valley fever. *Coccidioides* is endemic to hot, dry regions such as the Southwestern United States, Central America, and South America^1, 2^. The fungus grows in the soil as mycelia prior to disarticulating into the infectious arthroconidia spores. Upon aerosolization, spores are inhaled into the lungs where they become endosporulating spherules causing respiratory distress. 60% of Valley fever cases remain asymptomatic while 40% experience flu-like symptoms that mostly resolve on their own, and of these, 5-10% of infections result in chronic disease^1^. The biological factors contributing to acute or chronic coccidioidomycosis have yet to be fully elucidated. In addition, the disease is often misdiagnosed as bacterial pneumonia, resulting in 60-80% of these misdiagnosed patients being treated with several rounds of antibiotics prior to accurate diagnosis^3^. This is due to poor testing strategies and contributes to a growing problem not only of antibiotic resistance, but also antibiotic induced microbiome dysbiosis that contributes to several chronic disorders such as inflammatory bowel disease, rheumatoid arthritis, asthma, and type 2 diabetes, to name a few^4–7^. Antibiotic induced dysbiosis correlates to a prevalence of pathogenic bacteria^8, 9^. The use of antibiotics significantly shifts the lung microbiota repertoire resulting in less diversity and a higher abundance of resistant bacteria than in untreated lungs^10^. Increased susceptibility and colonization with *Salmonella*, *Shigella flexneri*, and *Clostridium difficile* in germ-free mice is associated with antibiotic treatment^11–14^. It is unknown if this shift in commensals correlates to a reduced ability to clear *Coccidioides* infection in coccidioidomycosis.

The microbiome utilizes multiple mechanisms of inhibition to protect against invading pathogens. Direct competition for host nutrients can inhibit pathogen colonization.^15^ However, to overcome competition, pathogens often use nutrients that are not preferred by resident gut bacteria. Host microbiome may also produce factors to protect their host niche from other bacteria, viruses, and fungi. These indirect mechanisms of protection involve promoting factors that enhance the intestinal epithelial barrier or promote innate and adaptive immunity to inhibit pathogen colonization^15, 16^. A soil *Bacillus subtilis-*like species displays antifungal activity against *Coccidioides* growth, with a clear zone of inhibition between fungi and bacteria when grown *in vitro*^17^. Whether host commensal bacteria can also inhibit *Coccidioides* by direct or indirect mechanisms is unknown. Furthermore, it is unknown how antibiotic treatment resulting from misdiagnosis further affects the interrelationship between the host lung microbiome and the invading fungal pathogen.

The 2007 Human Microbiome Project did not initially include the lungs as a site of investigation as it was long thought to be sterile^18^. Culture-dependent techniques posed a challenge in lung microbiome collection as microbial abundance is low compared to other sites of the body and only 1% of all bacteria are culturable in the laboratory ^18, 19^. 16s rRNA sequencing methods used to identify microbial communities in a healthy lung identified *Firmicutes*, *Bacteroidetes*, *Proteobacteria*, *Fusobacteria*, and *Actinobacteria* as the most prevalent families^18, 20^. At the operational taxonomic unit level, *Prevotella*, *Veillonella*, and *Streptococcus* are routinely identified as prevalent residents of the lung^20^. The lung is part of the lower respiratory system along with the trachea and primary bronchi. The upper respiratory tract consists of the nose, mouth, sinuses, pharynx, and larynx. Among healthy individuals, the microbiome of the upper and lower respiratory tracts are indistinguishable^21^. Recent studies of COVID-19 and respiratory syncytial virus (RSV) infections have explored differences between intestinal and respiratory microbiomes due to antibiotic treatment^22–25^. Until recently, most infection microbiome studies have focused on the influence of intestinal dysbiosis on infection^26^. It is unknown if the upper and lower respiratory tract or the intestinal microbiome change with infection of *Coccidioides* and influence *Coccidioides* growth. For the purposes of this study, we investigated the impact of cultured tracheal microbiota which we considered to be representative of the lung microbiota, on *Coccidioides* growth *in vitro*.

## Material and methods

### Mice

Six- to ten-week-old C57BL/6 male and female mice (JAX #000664, The Jackson Laboratories, Bar Harbor, ME, USA) were purchased or bred for experiments. Mice were housed and bred at the University of California Merced specific-pathogen free animal facility in compliance with the Department of Animal Research Services and approved by the Institutional Animal Care and Use Committee (protocol AUP21-0004). Mice from multiple dams were used for experiments.

### Agar plates

2x glucose yeast extract (GYE) agar plates were made in accordance to the following recipe: 2% w/v glucose (Fisher Scientific), 1% w/v yeast extract (Fisher Scientific), 1.5% bacteriological agar (VWR) in diH_2_O. GYE was autoclaved at 121°C for 1hr, poured into 100×15 mm^2^ petri dishes (Fisher Scientific) and stored at 4°C. Columbia colistin and nalidixic acid (CNA) agar with 5% sheep blood (5%SB-CNA) and chocolate agar medium agar plates were purchased from Fisher Scientific.

### Arthroconidia harvest

NR-166 avirulent *Coccidioides posadasii* (*Δcts2 / Δard1/ Δcts3)* derived from *C. posadasii* strain C735 was used for all experiments (BEI Resources, Manassas, VA, USA) ^27^. Fungal glycerol stock was inoculated into liquid GYE media and cultured for 3-7 days at 30°C, 150 rpm in a shaking incubator. Liquid culture was streaked onto GYE agar plates and grown for 4-6 weeks to reach confluency and appropriate desiccation. To harvest arthroconidia, fungi were scraped off the plate using cell scrapers into a conical tube in PBS. Collection was vortexed for 1 min prior to filtering through a 40 µm mesh filter to dislodge any arthroconidia withheld in the segmented mycelia encasing. Fungus was vortexed again for 1 min and washed twice with PBS (centrifuged at 12,000xg for 8 mins then 20 mins at room temperature (RT) with the break off. Fungal pellet was resuspended in PBS. Viability was assessed by plating 10-fold serial dilutions and colony counting 3-4 days post-plating. Arthroconidia suspension was stored at 4°C for up to 3 months. Complete protocol can be found in Mead et al. ^28^

### Tracheal and intestinal microbiota growth

The trachea and small intestine were harvested under sterile conditions by opening the chest cavity, cutting the trachea at the top of the bronchiole branching and base of the larynx. The trachea was cut in half, inverted onto a respective agar plate, and spread. 3-4 cm of the small intestine closest to the stomach was harvested, cleaned of fecal material and major mucus contents, cut in half, and spread onto a respective agar plate. Plates were incubated for 48 hrs at 30-37°C. If 80% confluency was reached from direct plating, plates were used for spike in inhibition assays. For the trachea, if ∼80% confluency was not obtained from direct plating, then tracheal microbiota was harvested from the plate into 2mL of PBS; serial dilutions were performed and plated for 48 hrs. The serial dilution from each trachea that yielded ∼80% confluency were used for spike in inhibition assays.

### 50/50 Inhibition assay

Small intestine was harvested as described above and spread across half the GYE plate. Blank and PBS spread plates were used as controls. Simultaneously, 50 arthroconidia were spread across the other half of the GYE plate. Plates were incubated at 30-35°C for 11 days.

### Spike-in inhibition assay

Trachea and small intestine were harvested and spread across the entire agar plate. Blank and PBS spread plates were used as controls. Plates were incubated for 48 hrs at 30-37°C prior to spiking in 50 arthroconidia on the edge of the plate. Plates were incubated for an additional 11 days at 30-35°C.

### Disk diffusion spike in inhibition assay

Trachea and small intestine were harvested and spread across GYE agar plates. Blank and PBS spread plates were used as controls. 100 µl of a broad-spectrum antibiotic cocktail (ampicillin, rifampicin, streptomycin, and neomycin; 50 µg/mL each) or PBS control were placed onto a 2 cm diameter Whattman paper circle disk and placed in the center of the host microbiota spread for 48 hrs at 30-35°C. After 48 hrs the disk was removed and 50 arthroconidia were spiked onto the center of the plate. Plates were incubated for an additional 11 days at 30-35°C.

### Inhibition measurements

Pictures of agar plates were taken on day 4, 7, and 11 and the area of *Coccidioides* growth was measured using ImageJ software (Wayne Rasband and contributors, Version 1.53k). The scale was set to 8.5 cm for every agar plate prior to tracing the area of *Coccidioides* colony growth. Area was determined based on these measurements and recorded.

### Whole organ harvest for 16s rRNA sequencing

Replicates were included from different dams and equal number of female and male mice were used for 16s rRNA sequencing experiments. Right lung lobes were harvested from male and female mice from several different dams and stored for bacterial extraction at −80°C.

### Bacterial DNA extraction

Tracheal and intestinal growth on agar plates were harvested using 1 mL of PBS via cell scraping, centrifuged for 10 mins at 7500 rpm. Microbial pellet was resuspended in 180 µl of enzymatic lysis buffer (20 mM TrisCl, pH8, 2 mM sodium EDTA, 1.2% TritonX-100, and 20 mg/ml lysozyme added immediately before use). Harvested right lung lobes were cut into small pieces and microbial content of all plated and whole organ samples were isolated using the DNeasy Blood and Tissue Kit (Qiagen) following manufacturer’s protocols for extraction of bacterial content. DNA concentration was determined using NanoDrop.

### 16s rRNA sequencing

16s rRNA sequencing was utilized to identify the bacterial abundance and composition of microbiota derived from right lung lobes and trachea and small intestine grown on different agar types. DNA extracts of bacterial samples were prepared according to the Illumina 16s Metagenomic Sequencing Library Preparation protocol. The Illumina protocol targeted variable 3 (V3) and V4 regions of the 16s ribosomal RNA gene for sequencing. PCR amplification of the target area was performed using the 2X KAPA HiFi Hot Start Ready Mix (070988935001, Roche). Reverse and forward amplicon PCR primers recommended by Illumina were used (16s Amplicon PCR Forward Primer= TCGTCGGCAGCGTCAGATGTGTATAAGAGACAGCCTACGGGNGGCWGCAG; 16s Amplicon PCR Reverse Primer= GTCTCGTGGGCTCGGAGATGTGTATAAGAGACAGGACTACHVGGGTATCTAATCC). After PCR amplification, the V3 and V4 amplicon were purified using AMPure XP beads (A63881, Beckman Coulter). To attach dual indexes and Illumina sequencing adapters, additional PCR amplification was conducted using the Nextera XT Index Kit (15032350, Illumina). The final library was purified once again using AMPure XP beads. Libraries were sequenced using Illumina MiSeq sequencer (Illumina).

### 16S rRNA analysis

All analyses were performed using R version 4.1.3 with DADA2 version 1.22.0. Sequence reads were first pre-processed to trim off the primer sequence and truncated at 245 bp length for forward reads and at 179 bp length for reverse reads, to facilitate the technical quality drop at the beginning of the forward reads and at the end of both forward and reverse reads, then processed through DADA2 pipeline to identify amplicon sequence variants (ASVs) with chimeras being removed. These ASVs were classified to the genus level using the Ribosomal Database Project naive Bayesian classifier in combination with the SILVA reference database version 138.1 with minBoot=50 (the minimum bootstrap confidence for assigning a taxonomic level).

Singletons were removed for all downstream analyses. We also removed ASVs with a phylum of NA and ASVs with ambiguous phylum annotation. Low-yield samples were not included in the downstream analysis (10-12 ng/µL or 6 ng/µL for whole organ lung samples). A total of 332 ASVs were identified in 25 samples, with 19 plated organ samples, 2 negative control samples (pooled blank 5%SB-CNA, chocolate, and GYE plate samples), 3 whole organ samples, and 1 positive control sample (ATCC 10 strain control). The smallest number of reads per sample is 24434 (the R lung lobes whole organ sample with ID 32). We further removed two plated trachea samples (IDs 5 and 20) as both their absolute abundance and relative abundance composition were significantly different from other samples in the same group (trachea samples plated on GYE plates for sample ID 5 and trachea samples plated on chocolate plates for sample ID 20). 31 ASVs were identified in those 2 negative control samples, and these ASVs were removed from trachea and intestine plated samples as well as the whole organ right lung lobe samples.

### Statistics

Preliminary inhibition experiments were used to perform power calculations in G* Power; T-test, Means: Difference between two independent means (two groups), A priori: compute required sample size, two tails, power= 0.90, α= 0.05, to define replicate requirements.

Two-way ANOVA statistical analysis was performed for the 50/50 inhibition assay, the intestine spike in inhibition assay on 5%SB-CNA and chocolate agar, and the trachea spike in inhibition assays on 5%SB-CNA and GYE agar data with Šídák corrections for multiple comparisons and a 95% confidence interval. Mixed effect analysis was performed for intestine spike in inhibition assays on GYE with Šídák corrections for multiple comparisons and a 95% confidence interval. Unpaired parametric t-test with Welch’s correction and a 95% confidence interval was performed on Day 7 disk diffusion assay.

## Results

Bacteria in the soil can exhibit an antagonistic effect on the growth of *Coccidioides* in vitro^17^. To determine if host microbiota has the potential to inhibit *Coccidioides* growth, we placed *Coccidioides* and host microbiota in direct competition *in vitro*. We began inhibition assay experiments with small intestine microbiota because the intestine has a dense bacterial population that grows well in vitro. This method allowed us to survey a broad and unbiased range of culturable host microbiota. By plating small intestinal microbiota and *Coccidioides* simultaneously on their respective halves of the agar plate, we provided an equal opportunity for the fungus and microbiota to compete for nutrients and space (Figure 1A-C). *Coccidioides* growth area was measured at days 4, 7, and 11 post-spread on control plates and compared to microbiota experimental plates. Although day 4 was not statistically significant, inhibition of fungal growth was observed by day 4 in the presence of the small intestine microbiota (Figure 1D). On day 7 and 11, *Coccidioides* growth area was significantly decreased when *Coccidioides* was placed in direct competition with the small intestine microbiota compared to controls. *Coccidioides* growth area averaged 45.5 cm^2^ on day 7 and 55.5 cm^2^ on day 11 in control plates whereas *Coccidioides* area averaged 31.8 cm^2^ on day 7 and 45.3 cm^2^ on day 11 against intestine microbiota. Thus, the small intestine microbiota has an antagonistic effect, inhibiting *Coccidioides* growth by 31.8%, 30.2%, 18.4% at day 4, day 7, and day 11, respectively.

**Figure 1:**
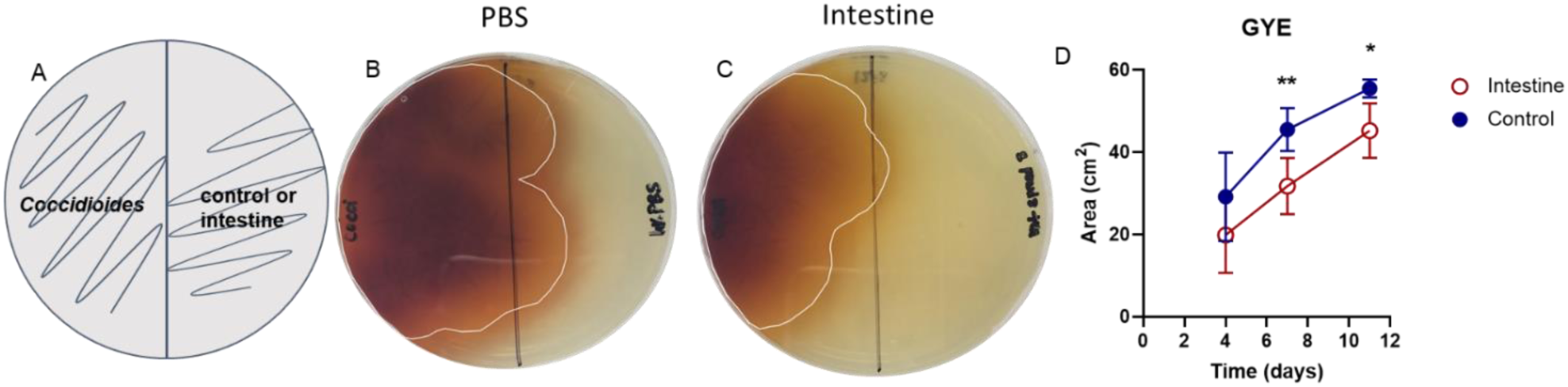
Intestinal mouse microbiota inhibits *Coccidioides* growth when in direct competition with *Coccidioides* on GYE agar in vitro. A) Experimental set up: 50 *Coccidioides posadasii* Δcts2/Δard1/Δcts3 arthroconidia were spread on half of a GYE agar plate and intestinal microbiota or PBS/Blank control was spread simultaneously on the other half of the plate. *Coccidioides* growth area was measured at day 4, 7, and 11. B) *Coccidioides* grown against controls (PBS/Blank) or C) in direct competition with intestinal microbiota. D) Area of *Coccidioides* growth at measured time points. Circles represent mean and errors the standard deviation; blue, closed circle=control, red, open circle=intestine; n=5-7. Performed mixed model statistical analysis; *p<0.05, ** p<0.005, *** p<0.0005.

While the direct inhibition assay served to assess the inhibitory potential of the host microbiota, we next sought to mimic the *in vivo* scenario in which the host microbiota is established prior to a *Coccidioides* infection. To achieve this, we allowed the small intestine microbiota growth to establish over 48 hours prior to spiking in *Coccidioides* to mimic an infection. GYE is the optimal growth media for *Coccidioides* whereas, 5%SB-CNA and chocolate agar media favor different bacterial communities and are used in clinical settings for diagnosis^29^. Thus, we used these media types to favor the growth of *Coccidioides* or host microbiota, respectively, to observe inhibitory potential in the presence of different nutrient sources. Small intestinal microbiota grown on GYE and 5%SB-CNA agar inhibited the growth of *Coccidioides*, which was depicted visually and numerically by decreased area of fungal growth on small intestinal microbiota plates compared to controls (Figure 2; Table 1). On GYE, *Coccidioides* grew to 31.2 cm^2^ on day 7 and 52.2 cm^2^ on day 11 in the controls as opposed to 24.3 cm^2^ on day 7 and 42 cm^2^ on day 11 when spiked onto an established lawn of small intestine microbiota (Figure 2D). On 5%SB-CNA, *Coccidioides* grew to 8 cm^2^ on day 7 and 8.9 cm^2^ on day 11 in controls as opposed to 4 cm^2^on day 7 and 4.8 cm^2^ on day 11 when spiked onto the established lawn of small intestine microbiota (Figure 2E). The small intestine microbiota selected for growth by chocolate agar media did not significantly inhibit fungal growth (Figure 2F). As expected, *Coccidioides* did not grow as well on 5%SB-CNA or chocolate agar compared to GYE; however, *Coccidioides* growth continued throughout the experiments.

**Figure 2:**
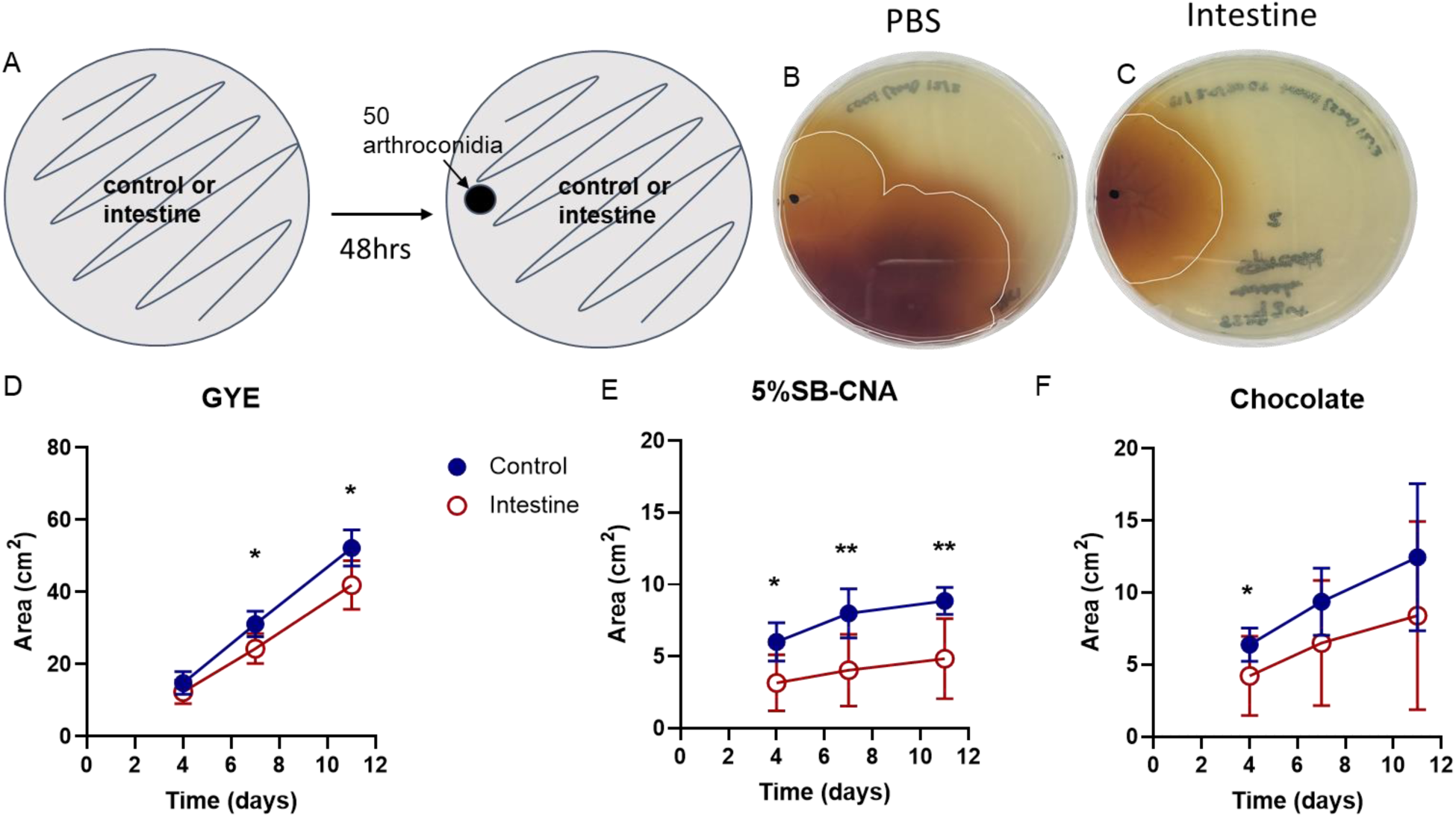
An established intestinal microbiota inhibits *Coccidioides* growth. A) Experimental set up: 50 *Coccidioides* arthroconidia were spiked onto a growing lawn of intestinal microbiota or control plate and the area of *Coccidioides* growth was measured at day 4, 7, and 11. B) *Coccidioides* spiked onto controls. C) *Coccidioides* spiked on a ∼80% confluently established lawn of intestinal microbiota. D-F) *Coccidioides* growth area at measured time points. Circles represent mean and errors the standard deviation; blue, closed circle=control, red, open circle=intestine on D) GYE, E) 5%SB-CNA, and F) and chocolate agar plates; n=7-17. Performed mixed model statistical analysis; *p<0.05, ** p<0.005, *** p<0.0005.

**Table 1:**
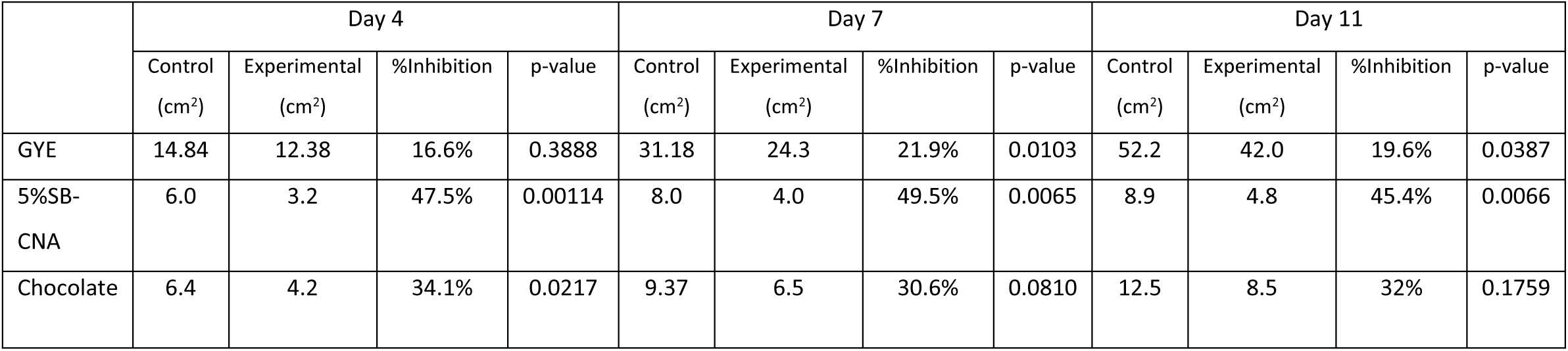
Intestinal spike in % inhibition.

|  | Day 4 |  |  |  | Day 7 |  |  |  | Day 11 |  |  |  |
| --- | --- | --- | --- | --- | --- | --- | --- | --- | --- | --- | --- | --- |
|  | Control (cm<sup>2</sup>) | Experimental (cm<sup>2</sup>) | %Inhibition | p-value | Control (cm<sup>2</sup>) | Experimental (cm<sup>2</sup>) | %Inhibition | p-value | Control (cm<sup>2</sup>) | Experimental (cm<sup>2</sup>) | %Inhibition | p-value |
| GYE | 14.84 | 12.38 | 16.6% | 0.3888 | 31.18 | 24.3 | 21.9% | 0.0103 | 52.2 | 42.0 | 19.6% | 0.0387 |
| 5%SB-CNA | 6.0 | 3.2 | 47.5% | 0.00114 | 8.0 | 4.0 | 49.5% | 0.0065 | 8.9 | 4.8 | 45.4% | 0.0066 |
| Chocolate | 6.4 | 4.2 | 34.1% | 0.0217 | 9.37 | 6.5 | 30.6% | 0.0810 | 12.5 | 8.5 | 32% | 0.1759 |

Since an established lawn of small intestine microbiota inhibited *Coccidioides* growth, we next assessed the inhibitory potential of host microbiota cultured from a more relevant organ for *Coccidioides* infection. Although the lung is the primary site of *Coccidioides* infection, lung microbiota is notoriously difficult to culture^30, 31^. Thus, we used the trachea as it is a part of the lower respiratory system and is indistinguishable from the upper respiratory system in healthy individuals^21^. Tracheal microbiota can also be cultured directly by trachea spread onto agar plates. GYE or 5%SB-CNA agar plates were used to favor *Coccidioides* or host microbiota, respectively. Tracheal microbiota did not grow to confluency on chocolate agar plates; thus these plates were not used. Tracheal microbiota cultured on 5%SB-CNA agar media displayed inhibitory potential on *Coccidioides* growth (Figure 3, Table 2). This inhibition was depicted visually and numerically by the decreased fungal growth area on plates with tracheal microbiota compared to controls. On GYE, *Coccidioides* growth showed differences in several individual experiments (Figure 3B,C), but was not statistically significant when the data was pooled (Figure 3D) perhaps due to inconsistent growth or low density of the inhibitory species. On 5%SB-CNA, *Coccidioides* grew to 7.7 cm^2^ on day 4 and 14 cm^2^ on day 7 in the controls as opposed to 3 cm^2^ on day 4 and 7.4 cm^2^ on day 7 when spiked onto an established lawn of tracheal microbiota (Figure 3E, Table 2). Thus, *Coccidioides* growth was inhibited to some extent by tracheal microbiota grown on both types of media.

**Figure 3:**
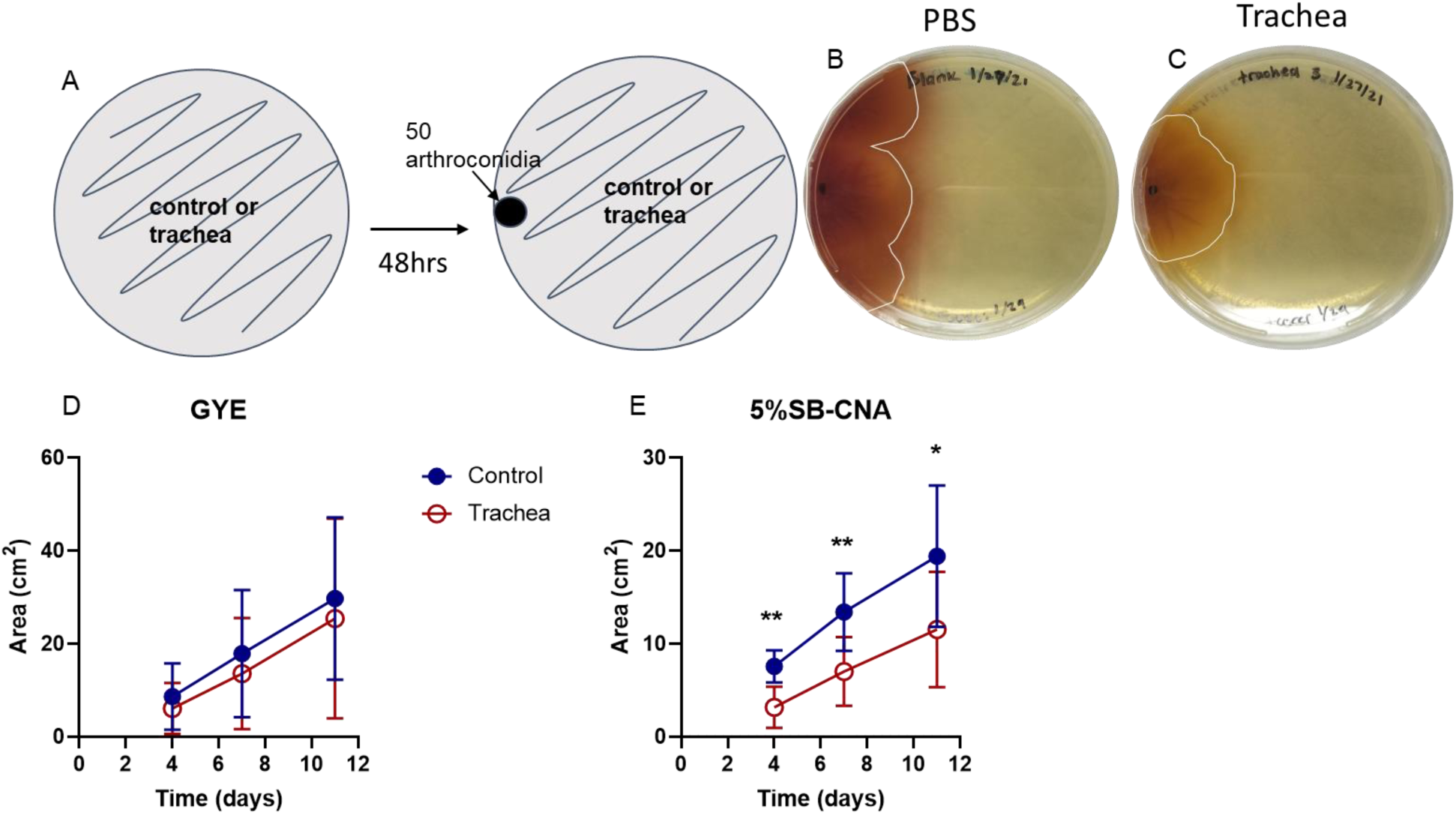
Tracheal mouse microbiota inhibits *Coccidioides* growth in vitro. A) Experimental set up: 50 *Coccidioides* arthroconidia were spiked onto a growing lawn of tracheal microbiota or control plate and the area of *Coccidioides* growth was measured at day 4, 7, and 11; B) *Coccidioides* spiked onto controls (1XPBS or blank); C) *Coccidioides* spiked onto a ∼80% confluent and established lawn of tracheal microbiota; D-E) Area of *Coccidioides* growth at measured time points. Circles represent mean and errors the standard deviation; blue, closed circle=control, red, open circle=trachea on GYE D) and 5%SB-CNA E) agar plates; n=7-15. Performed mixed model statistical analysis; *p<0.05, ** p<0.005, *** p<0.0005.

**Table 2:** Tracheal spike in % inhibition.

|  | Day 4 |  |  |  | Day 7 |  |  |  | Day 11 |  |  |  |
| --- | --- | --- | --- | --- | --- | --- | --- | --- | --- | --- | --- | --- |
|  | Control (cm<sup>2</sup>) | Experimental (cm<sup>2</sup>) | %Inhibition | p-value | Control (cm<sup>2</sup>) | Experimental (cm<sup>2</sup>) | %Inhibition | p-value | Control (cm<sup>2</sup>) | Experimental (cm<sup>2</sup>) | %Inhibition | p-value |
| GYE | 8.0 | 6.1 | 24.1% | 0.7734 | 17.9 | 13.6 | 24.1% | 0.8540 | 29.7 | 25.4 | 14.5% | 0.9539 |
| 5%SB-CNA | 7.7 | 3.0 | 61.3% | 0.0012 | 14.0 | 7.4 | 47.2% | 0.0051 | 20.1 | 12.0 | 40.3% | 0.0472 |

The Kirby-Bauer disk diffusion susceptibility test is typically used to determine the susceptibility of bacteria to an antimicrobial compound. Susceptibility is measured by the presence or absence of microbial growth around the disks. We used the disk diffusion assay to clear a zone of plated intestinal microbial growth using a cocktail of broad-spectrum antibiotics (ampicillin, rifampicin, streptomycin, and neomycin; 50ug/mL each), mimicking antibiotic treatment *in vivo*. PBS disks did not disrupt the surrounding bacterial growth and were used as a control. Fungal growth was not disrupted on control plates treated with disks soaked in PBS or antibiotics (Figure 4B). The area of growth between the two controls were not statistically significant, thus these data were pooled. When comparing *Coccidioides* growth on PBS versus antibiotic-treated host microbiota plates, we observe that the area of growth was larger on antibiotic disk-treated plates than on plates treated with a PBS disk (Figure 4C, D). Day 7 growth had the most pronounced differences with the area of Coccidioides growth being 6.25 cm^2^ on the host microbiota with the PBS disk versus 13.19 cm^2^ on the host microbiota with the antibiotic disks (Figure 4E). Thus, the elimination of the intestinal microbiota with the use of the antibiotic cocktail created a niche for *Coccidioides* growth. *Coccidioides* growth was inhibited when the microbiota was present, further confirming the potential of the microbiota to have an inhibitory effect on *Coccidioides*.

**Figure 4:**
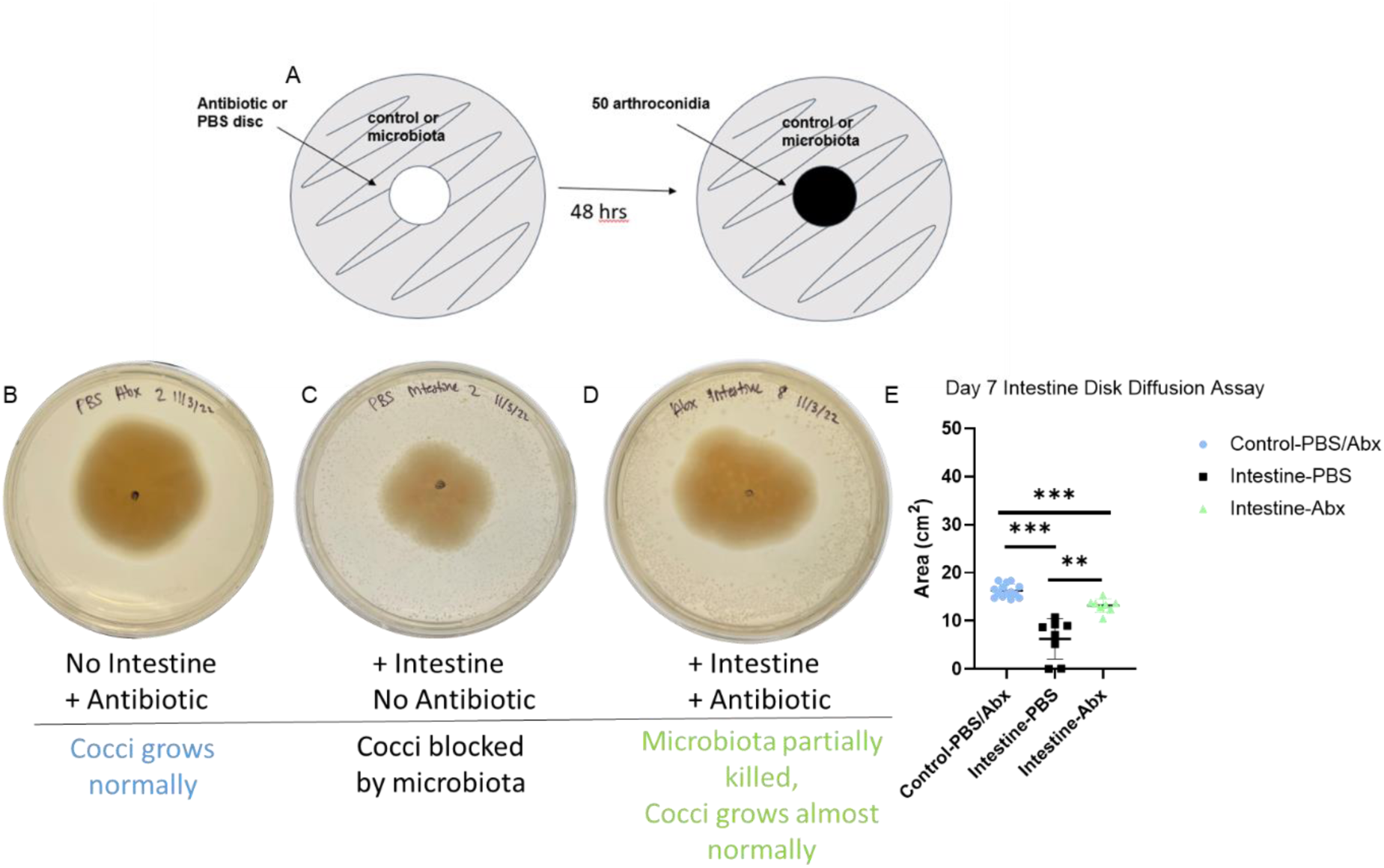
Antibiotic depletion of intestinal microbiota by antibiotic disk diffusion allows a niche for *Coccidioides* colonization and growth in vitro. A) Experimental set up: microbiota or control were spread, and an antibiotic or PBS control disk was placed in the center of the plate for 48 hrs. Disks were removed at 48 hrs and *Coccidioides* was spiked onto a growing microbiota lawn or control. *Coccidioides* growth area was measured at day 4,7, and 11; B) Representative pictures of *Coccidioides* spiked onto controls; or C) onto a growing microbiota lawn treated with PBS disk; or D) onto a growing microbiota lawn treated with an antibiotic cocktail (ampicillin, rifampicin, streptomycin, and neomycin; 50ug/mL each) disk; E) Area of *Coccidioides* growth at day 7. Errors present the standard deviation; blue, circles=control, black, squares= intestine spread with PBS disk, green, triangle=intestine spread with antibiotic disk) on GYE agar. n= 6-11. Performed mixed model statistical analysis; *p<0.05, ** p<0.005, *** p<0.0005.

To identify the bacteria responsible for inhibition of *Coccidioides*, 16s rRNA sequencing of tracheal and intestinal growth on GYE, 5%SB-CNA, and chocolate plates was performed. The absolute abundance of ASVs at the phylum level varied between replicates of each organ on each agar type (Supplemental Figure 1). However, relative abundance ratios of ASVs at the phylum level were fairly consistent among organ and agar type (Figure 5A). At the phylum level, the plated tracheal and intestinal growths were both primarily dominated by Firmicutes on all agar types and secondarily by Bacteroidota on 5%SB-CNA and chocolate agars (Figure 5A). On GYE, Proteobacteria was found on all trachea and intestinal replicates (Figure 5A). At the family level, plated tracheal growths were primarily dominated by *Staphylococcaceae* on all agar types, while plated intestinal growths were primarily dominated by *Lactobacillaceae* on GYE agar and *Staphylococcaceae* on chocolate agar (Supplemental Figure 2). 5%SB-CNA plates were dominated by different families among replicates (Supplemental Figure 2). Although the trachea and intestine are rather distinct in their environmental conditions and composition, bacterial composition was similar at the phylum level. This similarity diminished at the lower taxonomic levels, highlighting unique ASVs among the three agar plates (Figure 5B), although common ASVs did remain at the genus level. Bacteria from the *Staphylococcus* genus were shared between tracheal growths on GYE and 5%SB-CNA agar types that culture bacteria with inhibitory potential against *Coccidioides* in spike-in assays (Figure 5B, Table 3). One ASV from the family *Lactobacillaceae* was uniquely shared by the two agar types of interest, 5%SB-CNA and GYE (Figure 5B, Table 3). Since both the plated tracheal and intestinal growth showed inhibitory potential, we next sought to identify shared ASVs between the trachea and intestine samples plated on GYE plates and 5%SB-CNA plates, respectively (Figure 5C). Tracheal and intestinal growths on GYE shared 3 ASVs, as did growths on 5%SB-CNA (Figure 5C). On GYE plates, all three ASVs were of the *Lactobacillus* genus, while the 5%SB-CNA plates included ASVs from *Mitochondria*, *Lactobacillaceae*, and *Staphylococcaceae* families (Table 4). Comparing bacterial growth on agar types with inhibition of *Coccidioides* allowed further characterization of bacteria with inhibitory potential for future study.

**Figure 5:**
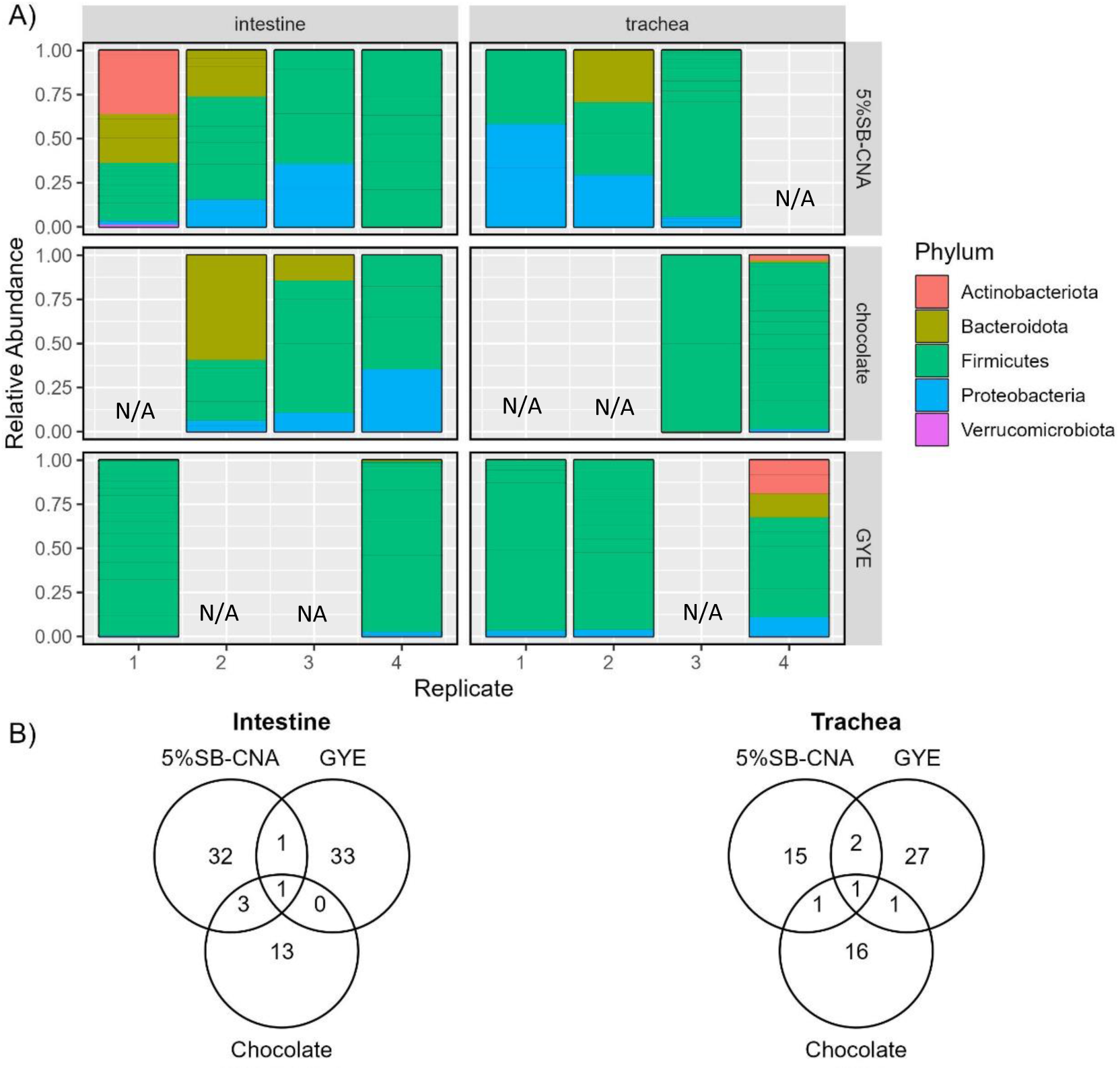
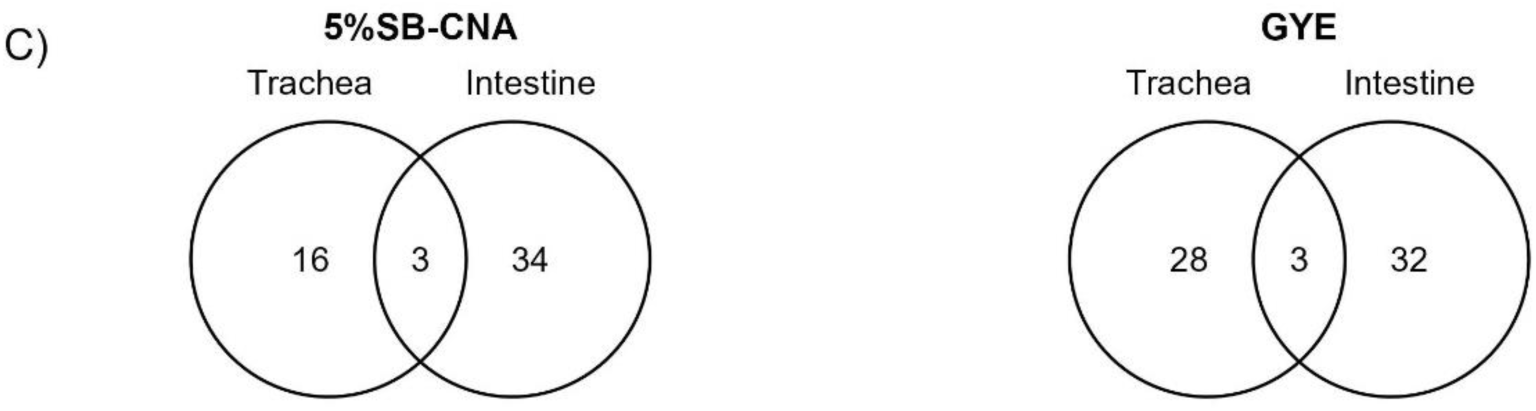
Bacterial composition and the relationships among tracheal and intestinal agar plates. A) Relative abundance of ASVs at the phylum-level in plated organ samples by plate type (row) and organ (column). B) Venn diagram depicting the number of shared and unique ASVs among three plates (5%SB-CNA, GYE, and chocolate) for the plated intestine samples (left) and trachea plated samples (right), respectively. C) Venn diagram showing the number of shared and unique ASVs between trachea and intestine for the samples plated on GYE plates (left) and 5%SB-CNA plates (right), respectively. Note: N/A= missing replicates removed due to low/poor DNA.

**Table 3:**
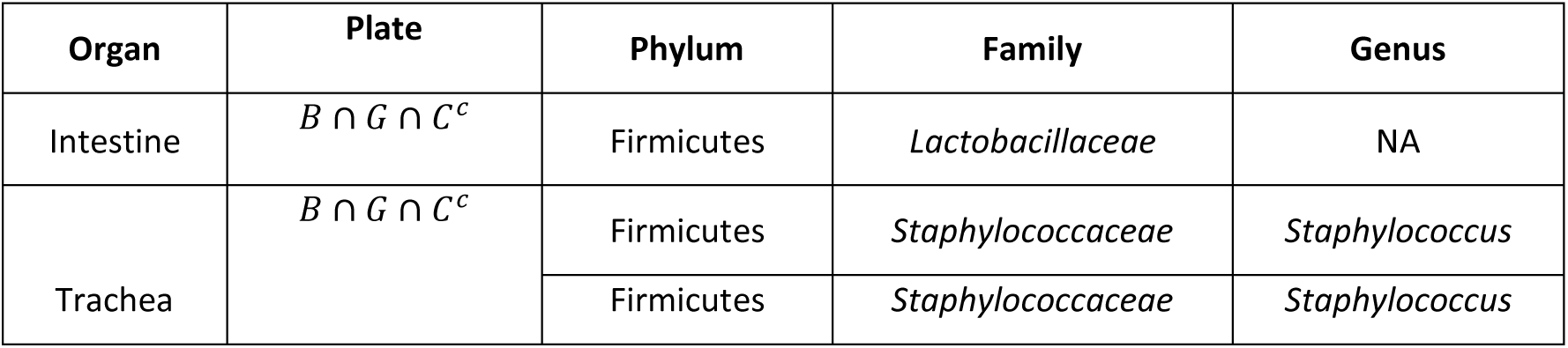
ASVs identified in the intestine or trachea samples when grown on 5%SB-CNA and GYE plates but not on chocolate agar. Note: B=5%SB-CNA, G= GYE, C= chocolate, c= excluding, ∩=shared, NA= could not be identified

**Table 4:** Information of ASVs present in both the trachea and intestine samples plated on GYE and 5%SB-CNA plates, respectively. Note: NA= could not be identified

| Plate | Phylum | Family | Genus |
| --- | --- | --- | --- |
| GYE | Firmicutes | *Lactobacillaceae* | *Lactobacillus* |
|  | Firmicutes | *Lactobacillaceae* | *Lactobacillus* |
|  | Firmicutes | *Lactobacillaceae* | *Lactobacillus* |
| 5%SB-CNA | Proteobacteria | *Mitochondria* | NA |
|  | Firmicutes | *Lactobacillaceae* | NA |
|  | Firmicutes | *Staphylococcaceae* | *Staphylococcus* |

As the lung microbiota is refractory to *in vitro* culture, mouse lung extracts were sequenced and compared to cultured intestine and trachea for overlapping bacterial identification. The lung microbiota was nearly evenly dominated by Proteobacteria, Firmicutes, and Bacteroidota in order of relative abundance at the phylum level (Figure 6A). Multiple ASVs were shared among the plated organs and right lung extracts. The trachea, lung, and intestine shared 2 ASVs, *Muribaculaceae* and *Bifidobacteriaceae* at the family level (Figure 6B). The trachea and lung shared *Achromobacter* at the genus level which is not present in the intestine. Lastly, the trachea and intestine predominantly shared bacteria in the *Lactobacillaceae* family which were not present in the lung (Figure 6B).

**Figure 6:**
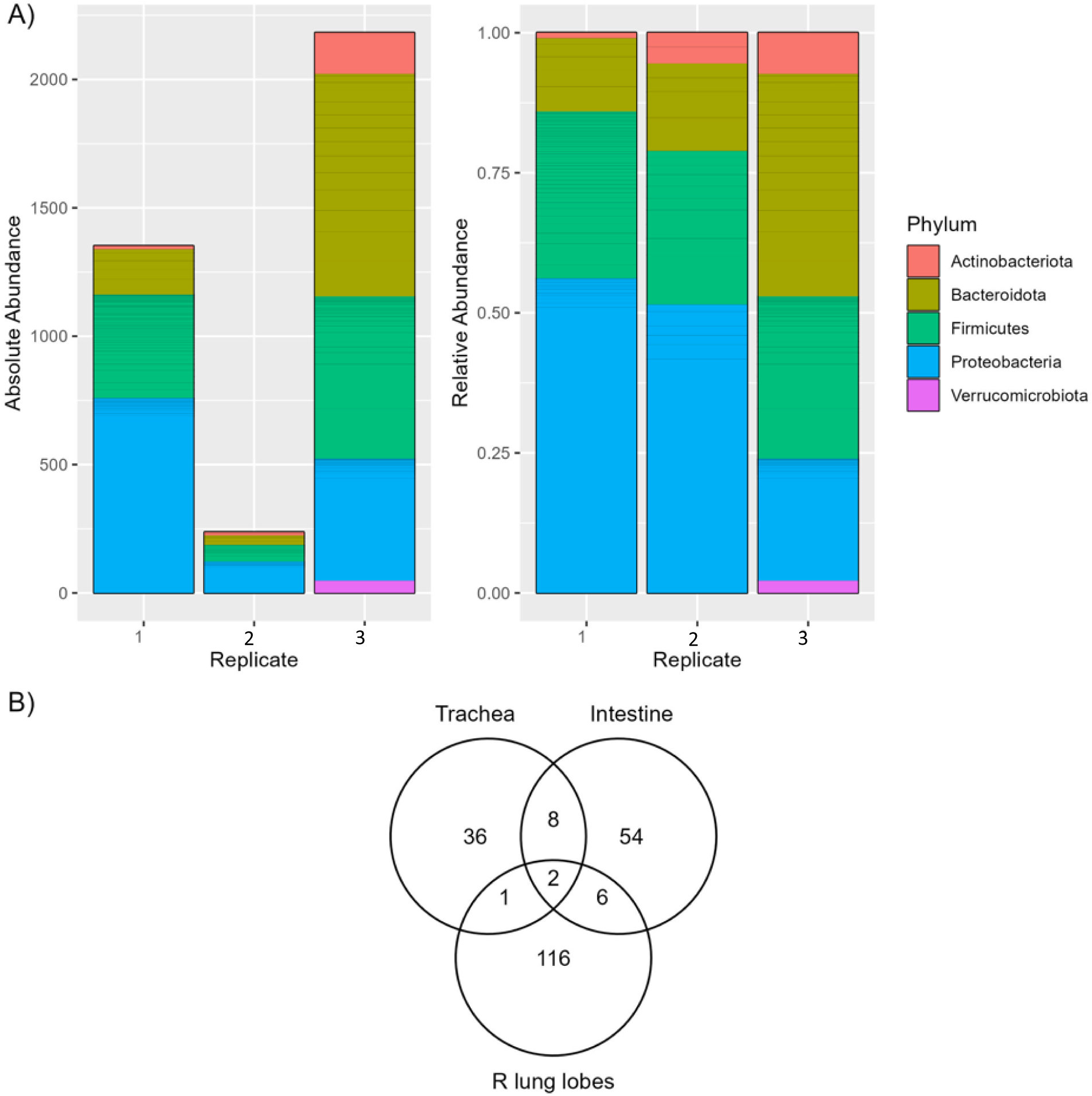
Bacterial composition of the right lung lobe. A) Phylum-level comparison of ASV absolute abundance (left) and relative abundance (right) in whole organ right lung lobe samples. B) Venn diagram showing the number of shared and unique ASVs between trachea and intestine for the samples on GYE and 5%SB-CNA plates and the whole right lung lobes. Note: R= right

## Discussion

The presence of host microbiota derived from either the intestine or trachea inhibits the growth of *Coccidioides in vitro*. The intestinal microbiota inhibits *Coccidioides* growth both when they are placed in direct competition and when the host microbiota is allowed to establish first. Thus, regardless of whether culture conditions provide an equal opportunity for the host microbiota and *Coccidioides* to compete or mimic an in vivo scenario in which we allow the microbiota to establish prior to infecting with *Coccidioides*, *Coccidioides* growth is inhibited. The tracheal microbiome is less dense in bacterial composition than the small intestine, thus not all plates spread with tracheal microbiota reached confluency. Therefore, only spike in inhibition assays were performed and only tracheal growths that reached ∼80% confluency were utilized. In these assays, tracheal microbiota also inhibited *Coccidioides* growth. There were differences observed in the level of inhibition based on the type of agar used. Microbiota cultured on 5%SB-CNA agar displayed inhibitory effects, whereas microbiota cultured on chocolate agar did not. The differences demonstrate that it is not simply the presence of microbiota that is responsible for inhibition, but rather the different type of microbes selected for by nutrients in the media type. 5%SB-CNA agar primarily selects for Gram positive bacteria, whereas chocolate agar primarily selects Gram negative bacteria but is relatively nonselective. *B. subitilis-* like species, a Gram positive bacteria species prevalent in the soil, displayed antifungal activity against *Coccidioides* in vitro.^17^ The bacteria identified in 5%SB-CNA agar should be considered for antifungal activity.

Due to misdiagnosis, 60-80% of Valley fever patients are treated with antibiotics^3^. To determine how perturbing an established microbiome would affect the inhibitory potential of the microbiota on *Coccidioides*, we depleted the microbiota with an antibiotic cocktail in vitro. Depletion of host microbiota through an antibiotic disk allowed a niche for *Coccidioides* growth. Although these are not in vivo studies, the *in vitro* data presented demonstrates the potential consequences of improper antibiotic treatment from misdiagnosing Valley fever patients with bacterial pneumonia and improper antibiotic treatment. Antibiotics may change the course of infection by altering the host microbiota and immune response^32, 33^. Antibiotic treatment can cause proximal changes in the microbial composition of the intestine that can lead to distal immunological changes in response to pulmonary infections. Antibiotics can also cause distal changes in the microbial composition of the lung. The in vitro data presented here demonstrate a direct influence of respiratory tract microbiota on *Coccidioides* growth.

Although the lungs are the primary site of *Coccidioides* infection, the trachea is also part of the lower respiratory system, and the intestine has proven to have influence on the respiratory infections through the gut-lung axis^23, 34, 35^. Thus, we sequenced bacterial growth from the trachea and intestine on the three agar types to identify levels of taxonomic order unique to the agar types that enabled growth of bacteria with inhibitory activity in our spike in assays. Cross comparison of the bacteria identified on each agar type and from each organ revealed shared ASVs among 5%SB-CNA and GYE agars, as well as those shared between the trachea and intestine samples plated on GYE and 5%SB-CNA. This allowed us to narrow down the candidates with potential inhibitory potential for future studies. To bring relevance to the pathogenesis of pulmonary *Coccidioides*, the right lung lobes of mice were sequenced for bacterial identification. As opposed to the plated cultures of tracheal and intestinal data, the lung data is from non-cultured whole lobe extracts due to the lung microbiota being notoriously difficult to culture^30, 31^. To avoid extensive manipulation of the resident microbiota, the whole lung was processed for extraction without culturing. We identified *Muribaculaceae* and *Bifidobacteriaceae* to be shared among all three organs and *Alcaligenaceae* to be shared between the trachea and right lung lobes (upper respiratory system) for future evaluation.

*Lactobacillus* and *Staphylococcus* were the predominant genus found in our plated sequencing data, likely because of abundance and ease of culturing these microorganisms. Previous work has demonstrated that the oropharyngeal, lung, and gut microbiota of healthy mice are dominated by *Lactobacillus* species^36^. However, it is possible that microorganisms that are challenging to culture have inhibitory potential on *Coccidioides* growth exist within the host microbiota but are difficult or virtually impossible to culture *in vitro*. Although this is a limitation of our study, *Lactobacillus* species have shown antifungal effects *in vitro* and been used as probiotics in viral and bacterial respiratory infection studies and improved infection outcomes^37–45^. Cell-free supernatants of *Lactobacillus plantarum* UM55 and *Lactobacillus buchneri* UTAD104 were tested against the fungal contaminant *Penicillium nordicum* and a reduction of radial growth and production of ochratoxin A were observed^46^. Acetic acid, indole lactic acid, and phenyllactic acid were the most effective in inhibiting *P*. *nordicum* growth and ochratoxin A^46^. *In vivo*, antibiotic induced dysbiosis during upper respiratory tract infection with influenza A virus is restored by *Lactobacillus casei* 431 and *Lactobacillus fermentum* PCC^38^. *Lactobacillus* strains restored the imbalance in the upper respiratory tract microbiome and re-upregulated pro-inflammatory cytokines^38^. Mice treated with heat-killed *Lactobacillus gasseri* TMC0356 were protected against influenza virus infection through stimulating protective immune responses^39^. Few *Staphylococcus* species have been used as probiotics for therapeutic treatment as most are opportunistic pathogens that cause disease. *Staphylococcus aureus* colonizes the nose and *Staphylococcus saprophyticus* colonizes the urinary tract. However, *Staphylococcus epidemidis* (*S. epidermidis)* has shown probiotic potential in multiple human and animal model studies. *S. epidermidis* has ameliorated infection by *Staphylococcus aureus*, *Moraxella catarrhalis*, Group A *Streptococcus*, influenza virus A, *Streptococcus pneumoniae*, and *Klebsiella pneumoniae*^47–52^. Treating mice with *S. epidermidis* NRS122 and streptomycin reduced colonization by *Staphylococcus aureus* BD02-31 compared to mice that received streptomycin alone^48^. In a mouse model of Influenza A, intranasally pre-colonizing with *S. epidermidis* limited the spread of influenza virus A to the lungs by modulating IFN-γ dependent innate immune mechanisms^52^. Yayurea A and B, small compounds isolated from *Staphylococcus delphini*, are expressed in a *Staphylococcus* species group^53^. These compounds have inhibitory potential against Gram negative bacteria^53^. Additionally, bacteriocins proteins produced by *S. epidermidis* inhibited *Micrococcus luteus*, *Corynebacterium pseudodiphteriticum*, *Dolosigranulum pigrum*, and *Moraxella catarrhalis*, bacterial species frequently found in human nasal microbiomes^54^. *S. epidermidis* bacteriocins might also be used against pathogenic bacteria. In addition to *S. epidermidis, Staphylococcus xylosus* VITURAJ10 also suppressed the growth of pathogenic strains of *Escherichia coli*, *Salmonella enterica*, and *Staphylococcus aureus*^55^. *Staphylococcus succinus* AAS2 also displayed antagonistic traits against *Staphylococcus aureus*^56^. These studies with other respiratory infections are evidence for the potential of *Staphylococcus* species to be used as probiotic treatment to improve infection outcomes.

Sequencing data from the lung as well as tracheal and intestinal plates revealed that *Bifidobacterium* was shared among the three organs (Table 5). Randomized, controlled human clinical trials and mouse models have proven the efficacy of using *Bifidobacterium* as a probiotic during respiratory tract infections, *Klebsiella pneumoniae*, influenza, and rhinovirus infection^57–63^. Oral treatment with commensal probiotic *B. longum* 5(1A) protected mice against *Klebsiella pneumoniae* pulmonary infection by activating Toll-like receptor signaling pathways that alter inflammatory immune responses^58^. A randomized controlled study also revealed that *B. animalis* subspecies lactis BI-04 affects the baseline of innate immunity in the nose^60^. Administering a single probiotic has resulted in amelioration of many pulmonary infections; however, probiotic cocktails have also proven to be effective. Administering *Lactobacillus rhamnosus* GG in combination with *B. longum* resulted in improved lung injury following experimental infection^64^. Thus, the bacteria with inhibitory potential against *Coccidioides* could be evaluated as probiotics alone or in combination for therapeutic treatment of coccidioidomycosis. This could provide a supplemental or alternative therapeutic to the existing antifungal therapies; however, further assessment would be necessary prior to implementation.

**Table 5:** Information of ASVs present in the trachea and intestine samples, plated on 5%SB-CNA and GYE plates, and whole organ right lung lobe samples. Note that there are four ASVs shared between intestine samples plated on 5%SB-CNA plates and trachea samples plated on GYE plates that are distinct from the ASVs shown in Table 4, they are denoted with * in this table. Note: T=trachea, I= intestine, L=lung, c= excluding, ∩=shared, NA= could not be identified

| Organ | Phylum | Family | Genus | NOTE |
| --- | --- | --- | --- | --- |
| $T \cap I \cap L$ | Bacteroidota | *Muribaculaceae* | NA | * |
|  | Actinobacteriota | *Bifidobacteriaceae* | *Bifidobacterium* | * |
| $T \cap I^c \cap L$ | Proteobacteria | *Alcaligenaceae* | *Achromobacter* | |
| $T \cap I \cap L^c$ | Firmicutes | *Lactobacillaceae* | *Lactobacillus* | |
|  | Firmicutes | *Lactobacillaceae* | *Lactobacillus* |  |
|  | Proteobacteria | *Mitochondria* | NA |  |
|  | Firmicutes | *Lactobacillaceae* | NA |  |
|  | Firmicutes | *Staphylococcaceae* | *Staphylococcus* |  |
|  | Firmicutes | *Lactobacillaceae* | *Lactobacillus* |  |
|  | Firmicutes | *Lactobacillaceae* | NA | * |
|  | Firmicutes | *Lactobacillaceae* | NA | * |

Among healthy individuals, the upper and lower respiratory tract appear indistinguishable^21^. However, the microbiota differs between the upper and lower respiratory tract and even within the lung among individuals with asthma, COPD, and cystic fibrosis^19, 65–67^. Recent studies on humans, macaque and mice revealed that viral and bacterial infections cause shifts in the landscape of lung microbiota^24, 68–71^. It is unknown whether *Coccidioides* infection causes microbiome shifts, nor how infection plus antibiotic treatment alters the lung microbiome. Our data suggests that an altered microbiome through antibiotic treatment may allow a niche for fungal growth. This is an area of study that requires further investigation in order to advise clinicians on the risks associated with antibiotic treatment during *Coccidioides* infection. Such findings could revolutionize the way infectious diseases are treated by leveraging microbiome interactions and probiotic therapeutics. Existing antifungal therapies for chronic and severe *Coccidioides* have unpleasant and severe side effects; exploring alternative treatments could improve patient outcomes and contribute significantly to our understanding of host-*Coccidioides* interactions.

---

**Supplemental Figure 1:**
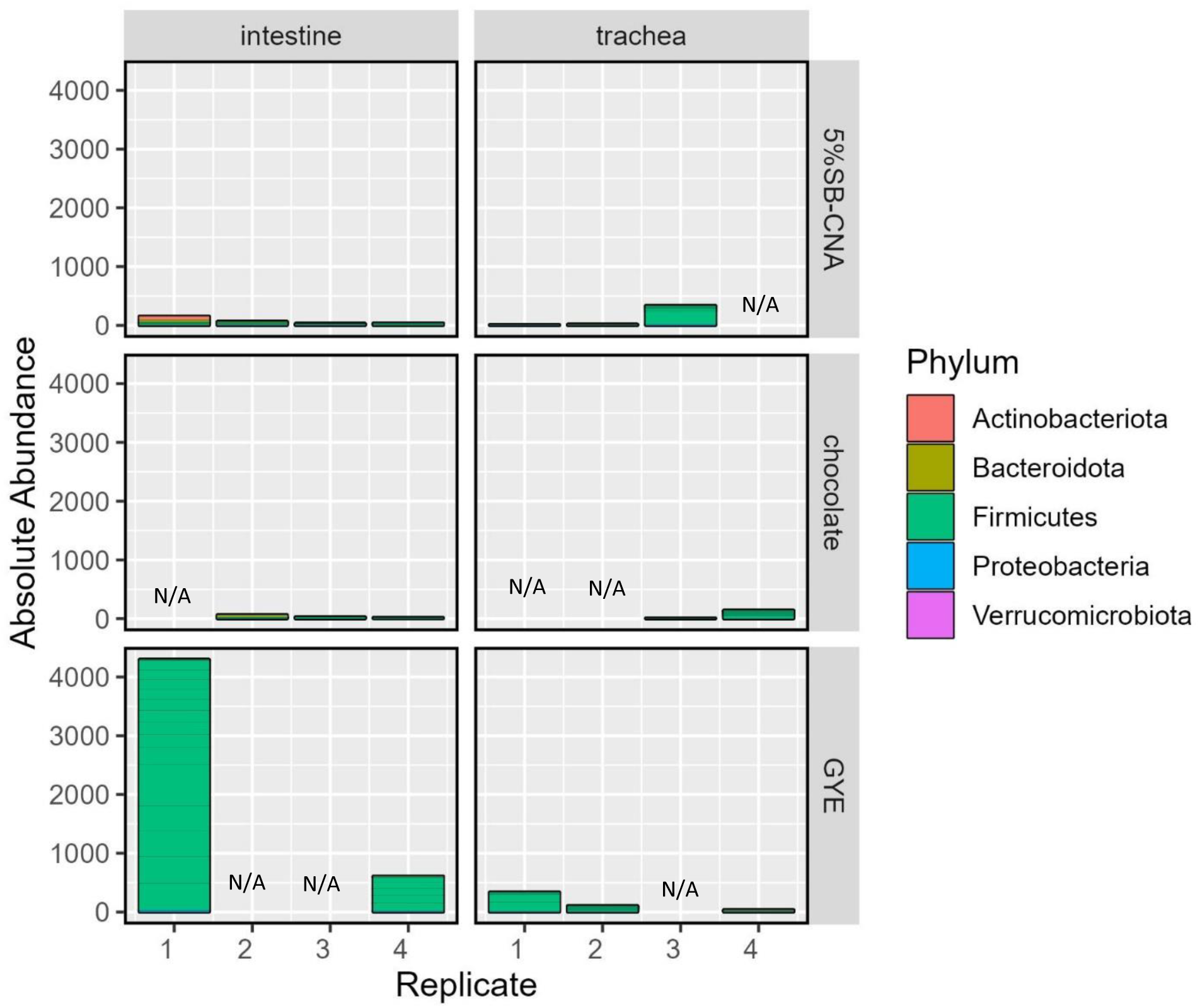
Phylum-level comparison of ASV absolute abundance in plated organ samples by plate type (row) and organ (column). Note: NA= missing replicates removed due to low/poor DNA.

**Supplemental Figure 2:**
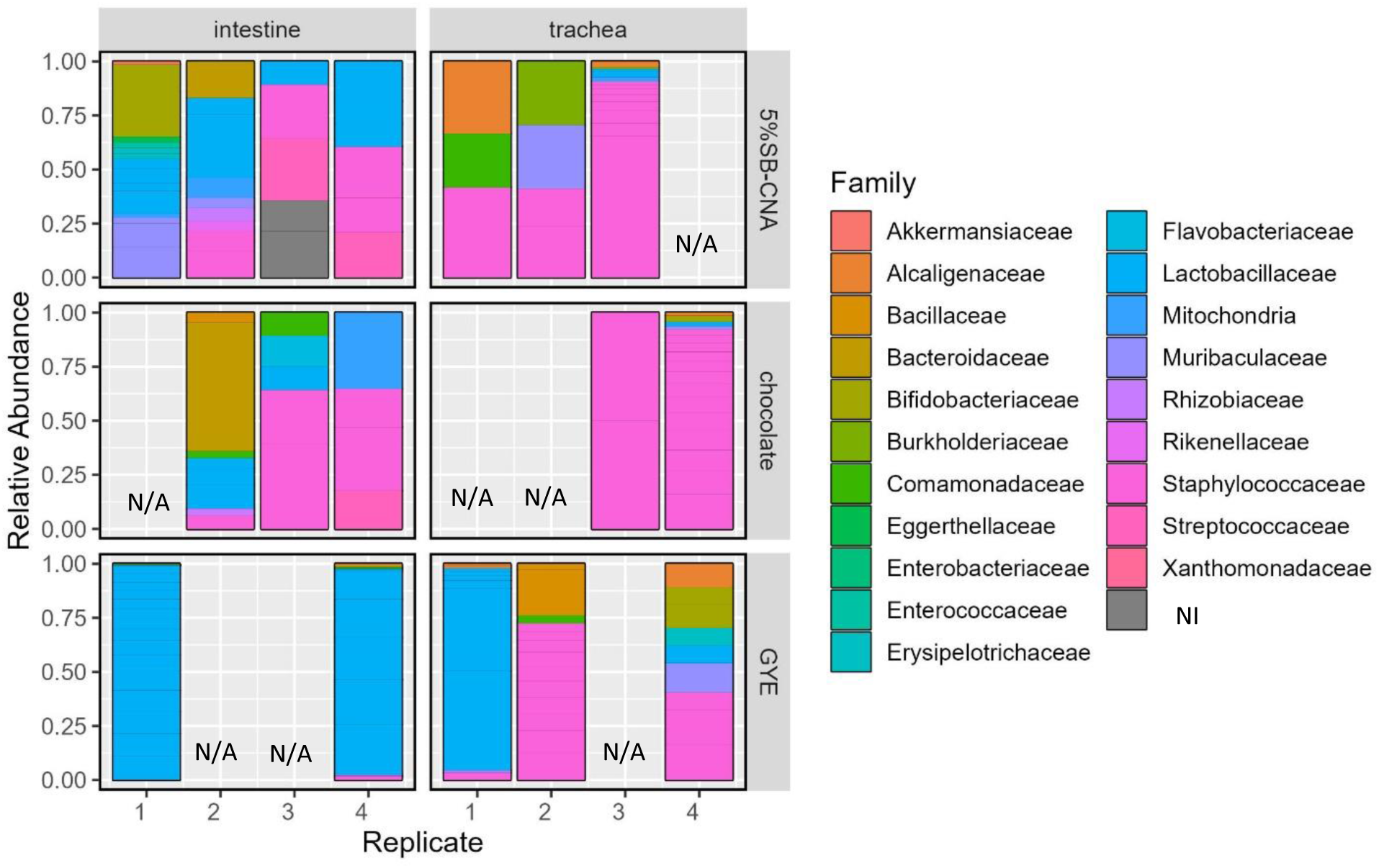
Family-level comparison of ASV relative abundance in plated organ samples by plate type (row) and organ (column). Note: NI= could not be identified, N/A= missing replicates removed due to low/poor DNA.

## Acknowledgments

The authors would like to thank Stem Cell Instrumentation Foundry-Tissue Culture (SCIF-TC) for tissue culture space, Health Sciences Research Institute for grant submission and management support, Hoyer lab members for experimental troubleshooting and conversations, *Coccidioides posadasii*, Δ*cts2*/Δ*ard1*/Δ*cts3*, NR-166 obtained from BEI resources, NIAID, NIH and Drs. Hernday and Nobile labs for conversations and shared equipment space. Portions of this work were performed under the auspices of the U.S. Department of Energy by Lawrence Livermore National Security, LLC, Lawrence Livermore National Laboratory under Contract DE-AC522-07NA27344.

